# Model for Deformation of Cells from External Electric Fields at or Near Resonant Frequencies

**DOI:** 10.1101/2020.05.01.072272

**Authors:** L. Martinez, A. Dhruv, L. Lin, E. Balaras, M. Keidar

## Abstract

This paper presents a numerical model to investigate the deformation of biological cells by applying external electric fields operating at or near cell resonant frequencies. Cells are represented as pseudo solids with high viscosity suspended in liquid media. The electric field source is an atmospheric plasma jet developed inhouse, for which the emitted energy distribution has been measured.

Viscoelastic response is resolved in the entire cell structure by solving a deformation matrix assuming an isotropic material with a prescribed modulus of elasticity. To investigate cell deformation at resonant frequencies, one mode of natural cell oscillation is considered in which the cell membrane is made to radially move about its eigenfrequency. An electromagnetic wave source interacts with the cell and induces oscillation and viscoelastic response. The source carries energy in the form of a distribution function which couples a range of oscillating frequencies with electric field amplitude.

Results show that cell response may be increased by the external electric field operating at or near resonance. In the elastic regime, response increases until a steady threshold value, and the structure moves as a damped oscillator. Generally, this response is a function of both frequency and magnitude of the source, with a maximum effect found at resonance. To understand the full effect of the source energy spectrum, the system is solved by considering five frequency-amplitude couplings. Results show that the total solution is a nonlinear combination of the individual solutions. Additionally, sources with different signal phases are simulated to determine the effect of initial conditions on the evolution of the system, and the result suggests that there may be multiple solutions within the same order of magnitude for elastic response and velocity. Cell rupture from electric stress may occur during application given a high energy source.

**Significance:** Cold atmospheric plasma jets (CAPJs) have been widely researched for their potential applications in cancer therapy. Existing research has focused mainly on the ability of CAPJs to deliver a mixture of reactive species which can be absorbed by cancer cells and induce cell death. The objective of our study is to investigate the mechanical effect of CAPJ electromagnetic (EM) waves on interacting cells. By coupling the EM waves associated with plasma frequency and cell viscoelastic response, we have developed a numerical tool to investigate cell damage either by mechanical or thermal loads. This work is motivated by the promise of EM waves to function as a sensitizing agent for cancer cells in preparation for chemotherapy.

## 1 Introduction

Experimental and numerical investigations have characterized both cell-generated and external electromagnetic (EM) fields acting on biological cells. Existing literature provides a basis for a numerical model presented in this paper, in which electrical stresses from cell-generated and external oscillating EM fields interact to deform cells. As summarized below, this interaction is often understood on the basis that biological cells behave as viscoelastic systems in which external forces induce a cell response.

Kositsky et al. (2001) summarizes experiments performed by Ukrainian and Russian scientists regarding the effect of high frequency electromagnetic radiation (EMR) at low thermal intensities on the human body.^1^ Among the important findings discussed in this report are the working principles that typically, human response to external EMR is resonant in nature, and that this response can be particularly observed in the milli-meter (GHz) EM wavelength regime. Identifying the target’s resonance frequency is a challenging aspect of tailoring the source EM devices to produce the desired effect. Blatt and Weiskopf (1952) proposed a numerical framework for the energy absorption cross-section of a resonant system from an EM source by means of an induced dipole.^2^ The authors also proposed that the absorbed power is a function of the incident (source) power and the absorption cross-sectional area. If the incident and absorbed power can be measured experimentally, then an estimation can be made regarding the cell’s energy absorption crosssection, and therefore its natural frequency. Escoffre et al. (2007) reviewed numerical and experimental studies examining the effect of EM fields on cell membranes, particularly the electropermeabilization of lipid membranes.^3^ For example, Rols (2006) performed an experiment to demonstrate permeabilization of hamster ovary cells by means of short high-voltage pulses. The author argued that when the applied external electric field reached values higher than the threshold dictated by the transmembrane potential, local transient permeable structures appear across the membrane, allowing for an exchange of hydrophilic molecules.^4^

As Tessie (2005) summarizes, the process of electropermeabilization can be described in five steps: (i) trigger (i.e., external field induces an increase in the transmembrane potential to threshold value in microsecond range); (ii) expansion (i.e., membrane time-dependent transition based on duration of electromechanical stress in millisecond range); (iii) stabilization (i.e., recovery of cell membrane organization when electric field falls below threshold value in millisecond range); (iv) resealing (i.e., minimization of transmembrane fluxes occurs in the seconds range); and (v) memory (i.e., cell behavior reverts to normal but some changes in the membrane properties recover in the range of hours). Riske and Dimova (2005) recorded the deformation and poration of unilamellar vesicles caused by an external electric field, for which the timescale of each step of deformation matches the descriptions summarized by Tessie (2005).^5^ Other studies have also reported on membrane permeabilization and the critical potential threshold necessary to achieve it^6,7,8,9,10^

Investigations have also sought to explain the basis of EM field generation within cells, and the role of these fields in regulating biological processes. Frolich (1968a, 1968b, 1969) theorized that biological cells may exhibit high frequency electrical oscillations (100 to 1000 GHz) as well as lower frequency oscillations within the radiofrequency regime, with the source of oscillations emanating from the cell membrane due to its high transmembrane potential.^11,12^ While this theory has been debated in the scientific community over the past decades,^13^ there is increasing evidence of cell-generated electrical oscillations. For example, some specialized cells are known to produce electrical oscillations via membrane depolarization and neuron firing.^14,15^ As Cifra (2015) postulates, an important follow-up question is whether and how biological cells in general produce EM fields.^16^ Cifra (2015) summarizes three distinct processes which produce cell-generated EM fields: (i) mechanical vibrations of polar structures such as membranes and cytoskeleton components (kHz-THz range); (ii) free ionic oscillations from chemical reactions (Hz-MHz range); and (iii) electronic oscillations such as DNA-driven conductivity (Hz-THz range). Understanding cell-generated EM field fluctuations may provide a foundation for expressing a cell’s natural oscillating pattern.

Experiments to characterize viscoelastic properties of various cells have also been performed. Faria et al. (2008) used atomic force microscopy to measure the elastic modulus of different prostate cancer cell lines. Their results reported a range of elastic modulus from 0 to 3000 Pascals.^17^ Guck et al. (2005) aimed to obtain cell deformation data and correlate it to cytoskeletal composition of MC-10, MCF-7, and modified MCF-7 human breast cancer cells.^18^ In the experiment, deformation was performed using a microfluid optical stretcher device composed of two diverging lasers which trap and then deform individual cells. The forces of deformation were reported to be 200 to 500 pico-Newtons. Optical deformability was measured by recording the change in position of axes at reference and deformation times, as well as by measuring the reference and deformed integrated stresses. The authors argued that optical deformability is a direct measure of the strength of the cytoskeleton from which time or frequency dependent complex shear moduli or other relevant properties can be obtained.

Suresh et al. (2007) performed a statistical summary of deformation results for the three cell lines investigated in Guck et al. (2005).^19^ The summary concluded that cancerous MCF 7 cells are more deformable than the normal MCF 10 (a ten percent increase in optical deformability), and that the metastatic modified MCF 7 cells are even more deformable than MCF 7 cells (another ten percent increase). Suresh et al. (2007) theorized that the increased deformability of cancerous MCF 7 and modified MCF 7 cells relative to normal MCF 10 was accompanied by a reduction in elastic rigidity. This behavior was likely due to the reduction in F-actin concentration by as much as 30 percent caused during the malignant transformation of the cell. The author also importantly summarized that eukaryotic cells contain three distinct molecules that serve as the structural elements in the cytoskeleton: actin microfilaments (elastic modulus ~ 1.3-2.5 GPa), intermediate filaments (1.5 GPa), and microtubulues (1.9 GPa). As such, building a cell cytoskeleton as a gel-like solid composed of a network of diverse biomolecules would account for an effective elastic modulus given by a contribution from each element.

Suresh et al. (2005) investigated the relationship between mechanical properties and subcellular structural organization in pancreatic cancer cells (panc-1). Cells were deformed by the microplate mechanical stretching method, in which the forces of deformation were as high as 300 nano-Newtons. The authors found that elastic response and energy dissipation was different between control panc-1 cells and panc-1 cells treated with SPC, a lipid that influences cancer metastasis. The effective spring constant was reduced from about 5 to 2 milli-Newtons per meter, due to the effect of SPC on normal panc-1 cells. Similarly, energy dissipation increased upon addition of SPC, signaling a lower cell response of SPC-treated cells. Cell imaging revealed a reorganization of a pan-cellular keratin network to a perinuclear configuration due to SPC treatment, which led the authors to argue that keratin appears to dominate deformability of the panc-1 cell.

Viscoelastic analytical models have been developed which can be used to investigate the mechanical response of cells to external stimuli. Fraldi et. al (2015) developed a numerical model to examine the frequency response of one-dimensional cells by considering three different viscoelastic models: (i) Voigt, (ii) Maxwell, and (iii) spring-pot based model.^20^ Yang et. al (2008) developed a two-dimensional method to predict the response of Dictyostelium cells under micropipette aspiration by employing a Maxwell viscoelastic model and analytical equations for the cell’s potential function, pressure, and velocity, as well as using a level-set function to define the cell structure.^21^ He and Qiao (2011) developed a computational fluid dynamics model for fluid-solid interactions which includes a deformation term that can be extended to solid or enclosed fluid structures which employ level-set functions to define the multiphase regimes as well as the Eulerian and Lagrangian grids.^22^

Based on the literature discussed above, it is evident that: (1) electromagnetic fields stress biological cells; (2) this stress can be resonant in nature; (3) cells have viscoelastic response (which may be a function of frequency and vary by cell type); and (4) permeabilization of the membrane occurs due to changes in the resting electric fields. The work outlined in this paper builds on this evidence to investigate how and to what extent cell deformation occurs. Specifically, the model presented in this paper assumes that a charged cell membrane receives mechanical loading via an electric stress from an external source. The cytoplasm and cytoskeleton of a cell act as a pseudo-solid with elastic properties. This pseudo-solid cell is suspended in a liquid medium and contains a natural vibration mode and frequency. Viscoelastic response of the cell is defined by the deformation matrix in the Navier-Stokes momentum equation. The energy absorbed by the cell is estimated by assuming that the energy contained in the EM energy source adds to the internal energy of the cell and that some of this energy is used in the viscoelastic response term. The extent of cell response is predicted as a function of time

## 2 Methods: Physical and Mathematical Model

The interaction between a source electric field and a solid cell-like structure is investigated to determine the extent of deformation, as well as to determine the magnitude of induced velocity and viscoelastic response. Figure 1 shows a diagram of this interaction, with the electric field acting on a cell suspended in a liquid medium. By altering the amplitude and frequency of the external electric field, deformation and viscoelastic response are solved for a cell with prescribed eigenfrequency and elastic modulus.

**Figure 1.**
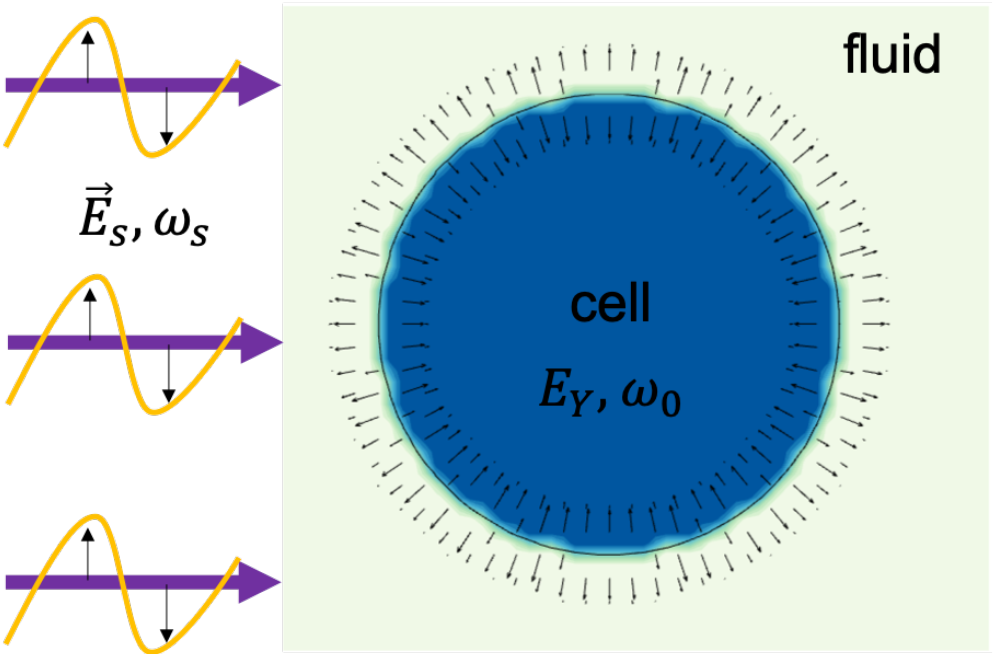
Cell natural vibration mode and external electric field. Cell eigenfrequency is assumed to be 28.4 GHz while the source frequency is assumed to be a distribution function with peak at the cell frequency.

The source-cell-fluid problem is solved by employing the Navier-Stokes equations, following fluid-structure interaction formulations.^23^ The momentum equation is solved for an incompressible fluid, and contains convective and viscous terms, as well as cell eigenfrequency force (natural oscillation), cell viscoelastic response to deformation, and the external electric field source. The Navier-Stokes continuity and momentum equations, in conservative form, are given by equations (2.1) and (2.2).

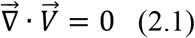

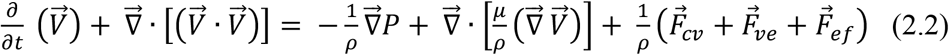

The system of equations are non-dimensionalized for computation in the in-house solver. These equations are found in Appendix A of supplementary materials. Existing literature on the bulk modulus of bilipid layered membranes shows a range of values (e.g., 10^7^ to 10^9^ Pascals) depending on the composition of the bilayer structure.^24,25^ This range of values provides the basis for the Reynolds number which characterizes the problem. In this study, the cell is made to oscillate harmonically and radially about its membrane with negligible damping during natural oscillations. The cell’s driving natural oscillation is given by equation (2.3) on a per unit length basis and can be interpreted as the cell’s eigenfrequency oscillating energy density.

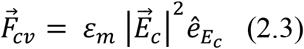

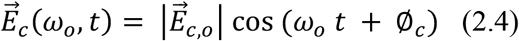

In equations 2.3 and 2.4, *ε_m_* is the membrane permittivity, 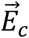 is the oscillating transmembrane electric field and 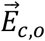 is its amplitude, *ω_o_* is the natural oscillating frequency of the cell membrane, and 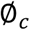 is the phase of the cell when the EM source reaches the cell initially. This natural oscillating frequency can be generally estimated based on dimensional analysis, i.e., by using the ratio of characteristic velocity of wave propagation in the medium (see equation 2.5) by the characteristic length of cell membrane (estimated to be in the order of nanometers).

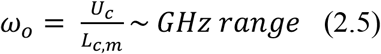

It is important to note that this approximation by dimensional analysis provides values of natural frequencies which include the operating range of frequencies created by atmospheric plasma devices. For example, a plasma jet with a density peak in the order of 10^19^ m^-3^ has been reported for in-house devices.^26^ This plasma density corresponds to an operating frequency of 28.4 GHz.

The viscoelastic response is formulated based on published works by He and Qiao (2011).^27^ The viscoelastic force is expressed according to equation (2.6).

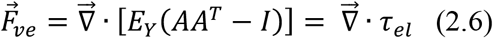

In equation (2.6), A is the deformation matrix, A^T^ is its transpose, and I is the identity matrix. The inverse of the deformation matrix for a two-dimensional structure is established. From this matrix, the corresponding deformation matrix and its transpose can also be obtained. These matrices are expressed by equation (2.7).:

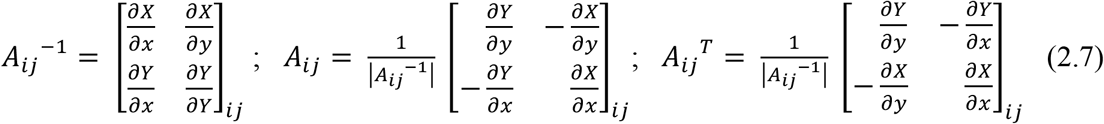

In equation (2.7), (X,x) are the (reference, current) current coordinates for a spatial grid node. The elastic modulus for biological systems such as cells may be anisotropic, and in the context of the formulation presented in this work, the modulus of elasticity would be a two-dimensional matrix for such anisotropic cases. The cases studied in Section 3 are formulated for an isotropic material with the same cytoskeleton modulus of elasticity for each degree of freedom.

The electric field source is made to oscillate about one reference dimension, which in our study is made to be the x-axis. The external stress driving the oscillating system can be expressed according to the Maxwell stress per unit length as shown in equation (2.8) and can be interpreted as the energy density delivered from the source.

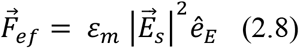

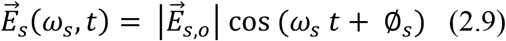

In equations (2.8) and (2.9), 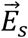 is the external electric field, 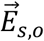 is its amplitude, *ω_s_* its frequency, and 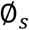 is the phase of the EM source as it reaches the cell. An energy equation in the Navier-Stokes formulation would include advective and diffusive terms as well as source terms accounting for addition and removal of energy to the system. For the purpose of this study the advective and diffusive terms are not examined since the necessary Prandtl number describing the biological cell has not been developed here. Instead, this model tracks the addition of potential energy from the EMF source and its conversion to kinetic energy in the form of cell viscoelastic response. The Maxwell stress is equivalent to the energy density absorbed by the system. Considering that any addition of energy results from this term, and loss occurs from kinetic energy expended during cell response, the change in internal energy (U) is given by equation (2.10).

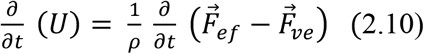

The electric field source used for the simulations presented in Section 3 is an atmospheric plasma jet, which produces a spectrum of electromagnetic wave energy densities at different frequencies. This energy spectrum can be represented by a peak function whose maximum is given by the plasma frequency, which is in turn a function of electron density. This means that different frequencies within a band of the plasma spectrum can deliver different magnitudes of electromechanical stresses. Thus, source electric field amplitude becomes another important parameter alongside frequency. If the plasma device is made to operate at or near cell resonant frequencies, electromechanical deformation may become significant. This would be accomplished by producing a plasma column with the required density such that the plasma frequency approaches the cell’s eigenfrequency.

Figure 2 shows preliminary experimental measurements of power density emitted by an inhouse atmospheric plasma jet. The results were obtained by employing a heterodyne method with an input signal ranging from 8 to 32 GHz. In these preliminary measurements, the maximum spatially averaged power density emitted from the device at steady state can be expected to be in the range from 0.05 to 0.15 mW per cm^2^ (or 0.5 to 1.5 W per m^2^), with higher values possible depending on the experimental parameters. Using equation 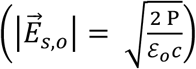 for the contribution of an electric field to power in an EM source, the average electric field can be found from the measured power density. This calculation yields a possible range of electric field amplitudes roughly between 10 to 100 V/m. Because the power measurements are spatially averaged, the maximum local value of power (and electric field) may be much larger. For example, consider experimental results which show that the electric potential value of the ionization wave in the plasma jet above a water target has been measured to be about 1.8 kV.^28^ In this experiment, the ionization wave size was measured to be in the millimeter range, and its average propagation velocity was estimated to be 9.3 km/s. Then, the static electric field of the IW is calculated to be about 1000 kV/m, with a power density of 82 kW per m^2^. These calculations are useful in determining a wide range of power delivered to the cell. While the source power is important in determining the stress that will be imposed on the cell, the source frequency is the critical factor in determining whether the cell experiences a stress at all. Ample documentation is available for the operating parameters and measurements techniques for the atmospheric plasma jets developed inhouse.^29,30,31^ Using these experimental results as references, an EM source function can be developed for the numerical model. Figure 3 shows the energy density of the EM source as a function of frequency used in the model, following a Gaussian distribution with a full width at half maximum (FWHM) of 5 GHz.

**Figure 2.**
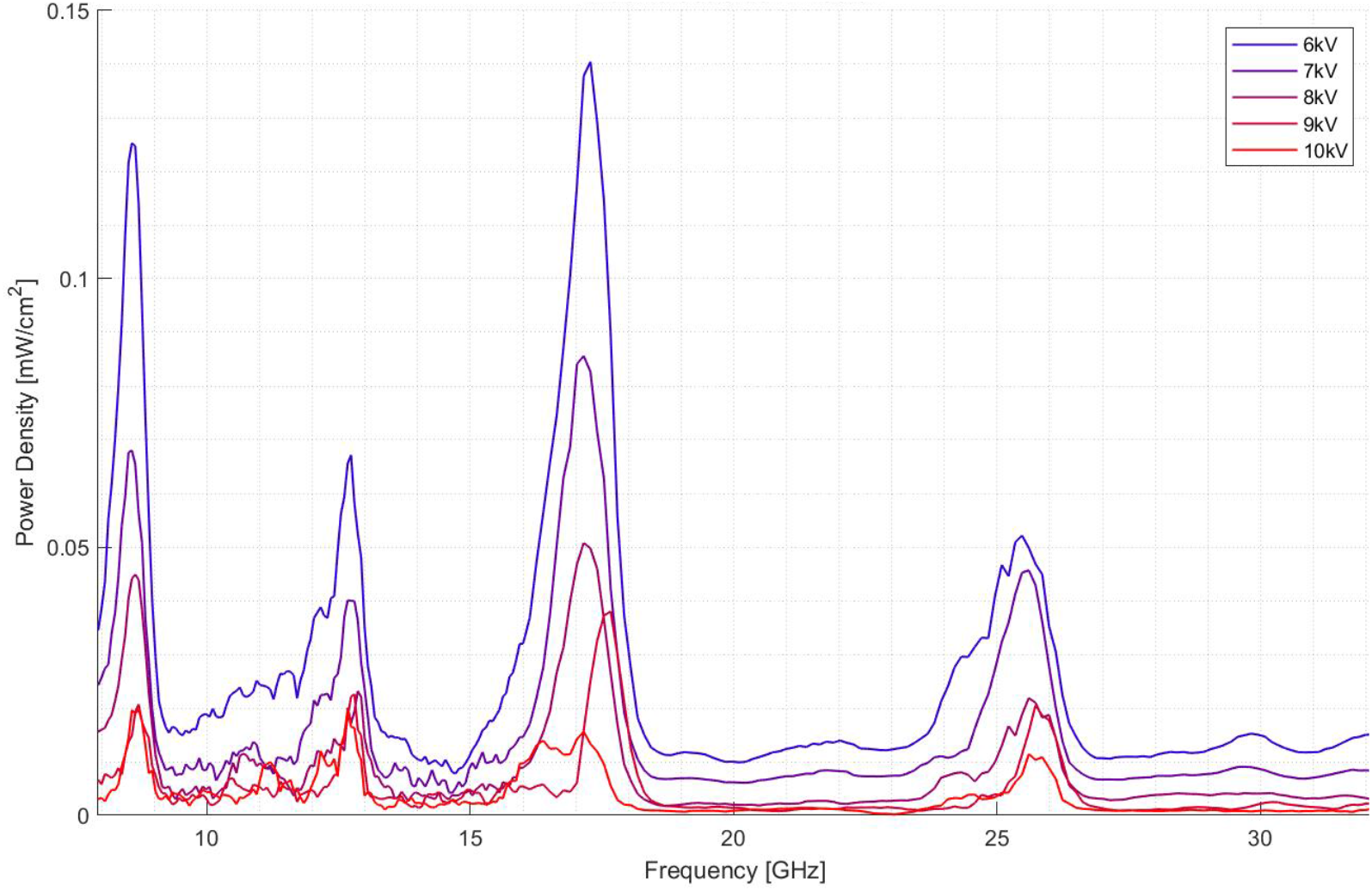
Power density measurement of an inhouse atmospheric plasma jet.

**Figure 3.**
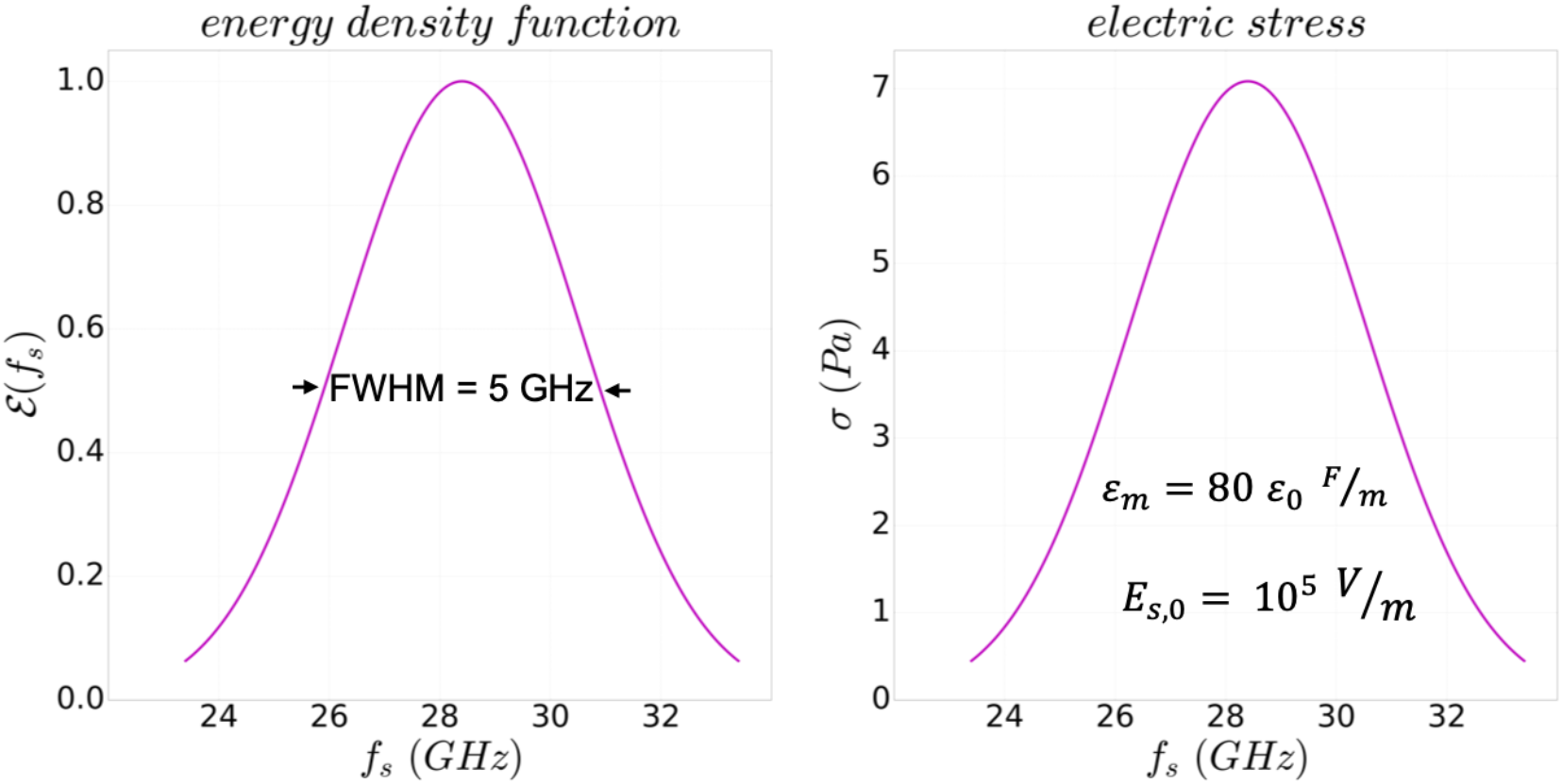
Energy density of jet with FWHM = 5 GHz, and corresponding distribution of electric stress corresponding to a peak electric field of 100 kV/m.

For the purpose of this study, dimensional analysis is used to estimate the eigenfrequency of a biological cell that would be treated with the EM source. Using reported values of plasma density and operating frequencies of in-house atmospheric plasma devices, the characteristic frequency is estimated to be about 28.4 GHz. This value falls within the range of oscillating frequencies for biological structures, and within the range of identified peaks of spatially averaged power (see Figure 2).^32^ Generally, if the target frequency is known, the plasma device may be tailored to provide a near resonant frequency. For example, if the frequency of a biological structure oscillating at 1 GHz matches the plasma frequency 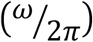, the corresponding plasma electron density would need to be in the order of 10^16^ m^-3^, which is a density also reported for in-house devices.^33^ For a plasma frequency of 28.4 GHz, the corresponding electron density would be in the order of 10^18^ m^-3^, which has also been reported in previous studies.^34^ The amplitude of the oscillating electric field across the cell membrane is based on a subthreshold membrane potential oscillation of 1 mV.^35^ The electric field amplitude is then calculated assuming a membrane thickness of 10 nanometers, which yields and electric field of 100 kV/m.

The intensity of oscillation is characterized by a formulation based on a driven (i.e., electric field) and damped (i.e., cell response) harmonic oscillator. This approach is justified by considering that the membrane contains a bound charge density and that cell response and oscillation velocity may reach steady state. The intensity of oscillation is given by equation (2.11) and its derivation is provided in Appendix B of supplementary materials.

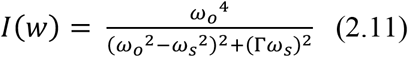

In equation (2.12), *ω_o_* is the natural oscillating frequency of the membrane, *ω_s_* is the source electric field frequency, and Γ is the damping coefficient. The damping coefficient is constructed from the structure’s amplitude and frequency of motion. The cell’s elastic response provides for the amplitude and the oscillating velocity gives the frequency. Cell oscillation and deformation are captured through implicit advection of the level-set function set which has been applied for multiphase flows.^36,37,38,39,40,41^ The level-set function is a scalar function that approaches zero at the cell-liquid boundary and distinguishes between the cell and host liquid medium by storing signed normal distances from the boundary. Distances are positive in phase one (cell) and negative in the other (host medium). The topology of the cell membrane is modified by solving a level-set advection equation given equation (2.12).

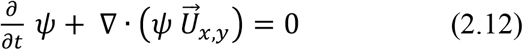

In equation (2.12), *ψ* represents the signed distance function. After advection, a selective redistancing technique is applied to preserve the membrane topology during each time step. ^42^ This re-distancing step is necessary because equation (2.12) is not inherently mass conservative, and if used by itself can lead to artificial changes in the membrane topology by not preserving the sharp gradients in the signed distance function. This step is numerical in nature, as it does not change the result from equation (2.12). The authors have previously employed this technique to analyze plasma contact spots on a gaseous-aqueous interface layer^43^, and has been validated elsewhere^44^. The re-distancing equation (2.13) is thus solved through multiple iterations in pseudo-time *τ* to satisfy mass conservation and preserve the signed nature of the distance function.

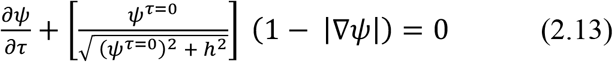

In equation (2.13), *h* is the grid spacing. The numerical framework presented in this section is implemented within our in-house multi-physics solver which has been validated for numerical accuracy and performance.^45,46^ The Navier-Stokes equations are solved on a block-structured AMR grid using an explicit fractional step method. For pressure, a hybrid multigrid-HYPRE Poisson solver is used. The level-set implementation within the solver has also been validated and provides an efficient way to track phase interface dynamics.^47^

## 3 Results and Discussion

### 3.1 Effect of source frequency-amplitude couplings on cell response

This section presents results on parametric studies for source amplitude and frequency effects on cell deformation. To investigate the near resonant effect, five source frequency and energy density amplitude combinations were examined. Table 1 summarizes these trials.

**Table 1.** EMF source energy and frequency trials

| **Table 1. EMF source energy and frequency trials** |  |  |  |
| --- | --- | --- | --- |
| **$\omega_0$ (GHz)** | **$\omega_s/\omega_0$** | **$E_s$ (N/C)** | **$S_s$ (Pa)** |
| 28.4 | 0.95 | 8.0e4 | 4.54 |
|  | 1.05 | 8.0e4 | 4.54 |
|  | 1.0 | 1.0e5 | 7.08 |
|  | 0.9 | 4.1e4 | 1.19 |
|  | 1.1 | 4.1e4 | 1.19 |
| Table 1. Trials corresponding to coupled energy-frequency effect on deformation. |  |  |  |

The coupled set of equations governing the dynamics of the problem are solved in non-dimensional form. The characteristic velocity of the system corresponds to the propagation of pressure waves emanating from the membrane as it is stressed. Thus, velocity is given in equation 3.1 by using the bulk modulus (*K_v,m_*) and density (*ρ_m_*) of the membrane.

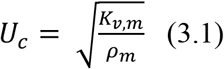

Adopting an effective elastic modulus of 10^7^ Pascals and density of 10^3^ kg/m^3^, the resulting characteristic velocity is 100 m/s. The elastic modulus was not chosen to represent a specific biological cell. However, it falls within the reported range of values for bilipid layered membranes, as discussed in Section 2. The characteristic length scale is chosen to be 10 micrometers, which is the diameter of the structure. Two non-dimensional numbers result from non-dimensionalization of the momentum equation; the Reynolds number and a number labeled here as the frequency number (Nf). Equation 3.2 shows this frequency number, which couples fluid characteristic time scale with the frequency of resonance.

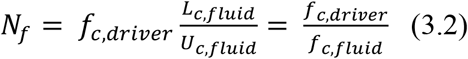

The frequency number is defined in the domain Nf ≥1, since the implication of a value less than one is that the cell has responded to a stress cycle before it has occurred. The physical implication of Nf > 1 is that the time to complete one characteristic driving stress cycle is smaller than the time it would take for the cell to fully respond to that cycle. Then, a value of Nf = 1 implies an instantaneous response of the entire structure. This is a significant parameter, as it provides insight into how sensitive a structure is in responding to frequency-dependent applied forces. Table 2 summarizes all of the characteristic values necessary to make the momentum equation non-dimensional. The derivation is provided in supplementary Appendix A.

**Table 2.**
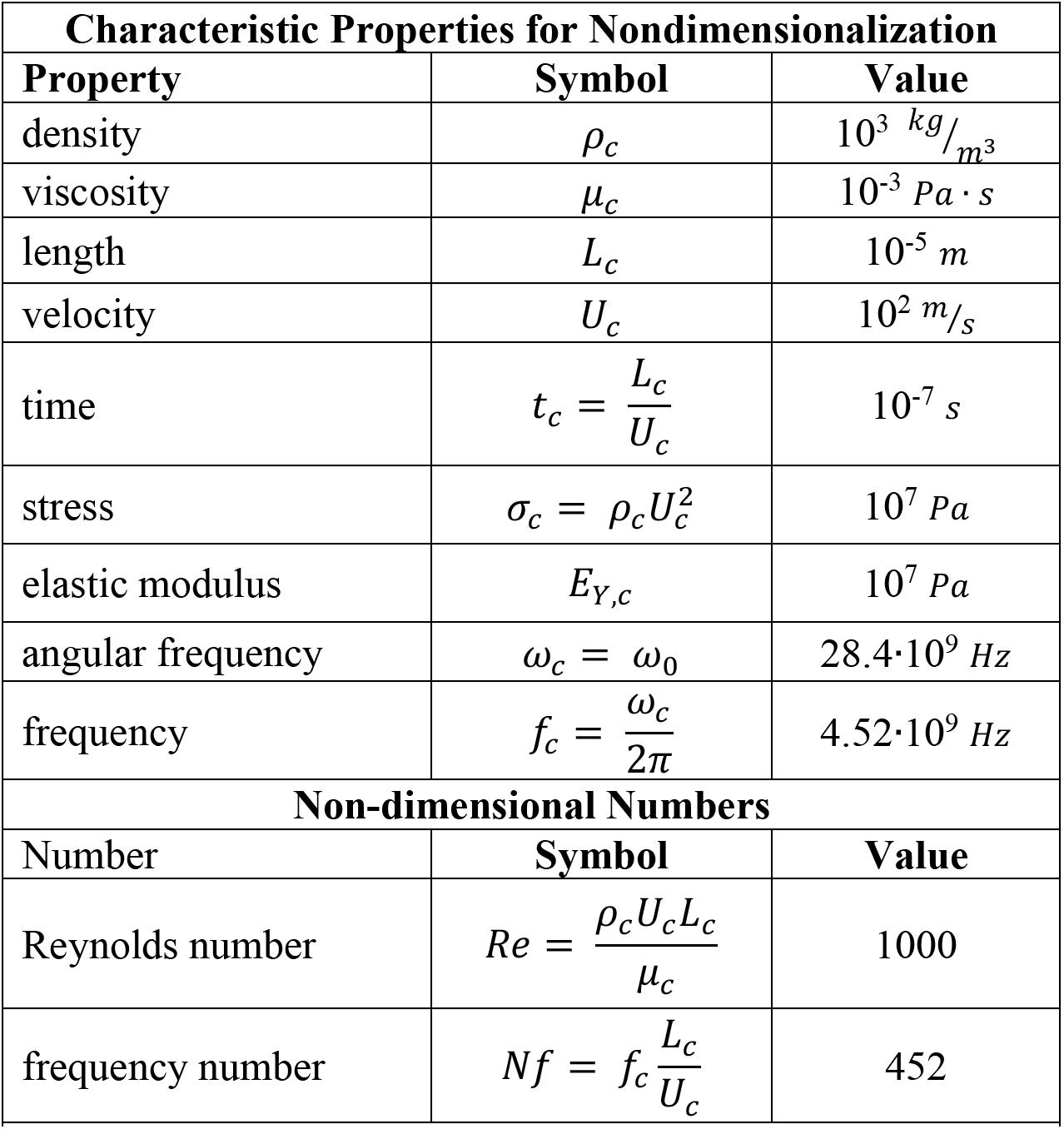
Characteristic values and non-dimensional numbers.

| **Characteristic Properties for Nondimensionalization** |  |  |
| --- | --- | --- |
| **Property** | **Symbol** | **Value** |
| density | $\rho_c$ | $10^3 \text{ kg}/\text{m}^3$ |
| viscosity | $\mu_c$ | $10^{-3} \text{ Pa} \cdot \text{s}$ |
| length | $L_c$ | $10^{-5} \text{ m}$ |
| velocity | $U_c$ | $10^2 \text{ m}/\text{s}$ |
| time | $t_c = \frac{L_c}{U_c}$ | $10^{-7} \text{ s}$ |
| stress | $\sigma_c = \rho_c U_c^2$ | $10^7 \text{ Pa}$ |
| elastic modulus | $E_{Y,c}$ | $10^7 \text{ Pa}$ |
| angular frequency | $\omega_c = \omega_0$ | $28.4 \cdot 10^9 \text{ Hz}$ |
| frequency | $f_c = \frac{\omega_c}{2\pi}$ | $4.52 \cdot 10^9 \text{ Hz}$ |
| **Non-dimensional Numbers** |  |  |
| **Number** | **Symbol** | **Value** |
| Reynolds number | $Re = \frac{\rho_c U_c L_c}{\mu_c}$ | 1000 |
| frequency number | $Nf = f_c \frac{L_c}{U_c}$ | 452 |
| Table 2. Characteristic values and non-dimensional numbers. |  |  |

As the cell experiences deformation from the EMF source, all frequency-amplitude couplings within the energy spectrum will affect the cell. Here, the couplings are solved individually and in aggregate in the NS equation to investigate the effect of each and compare to the total effect of the energy spectrum. Figure 4 shows that the total effect is greater than the individual couplings combined, and therefore cell elastic response is not a linear solution. The evolution of total elastic response exhibits a magnified unsteady period due to the temporal fluctuations of individual couplings (ωs, Es). The result also confirms that for individual couplings, maximum cell response is achieved at the resonant frequency-energy coupling. At steady state the contributions from frequencies with equal intervals from resonance produce nearly the same maximum value in elastic response. At cell response steady state, the structure oscillation properties such as velocity, damping, and intensity can be observed to follow cyclical pattern based on the driving external stress. Figure 5 shows one cycle of electric stress and corresponding induced maximum cell velocity. Oscillating velocity for the total effect is consistent with the observation that the total solution is nonlinear relative to the individual solutions. Viscoelastic response steady state was achieved after 3,285 stress cycles in 1.15 microseconds.

**Figure 4.**
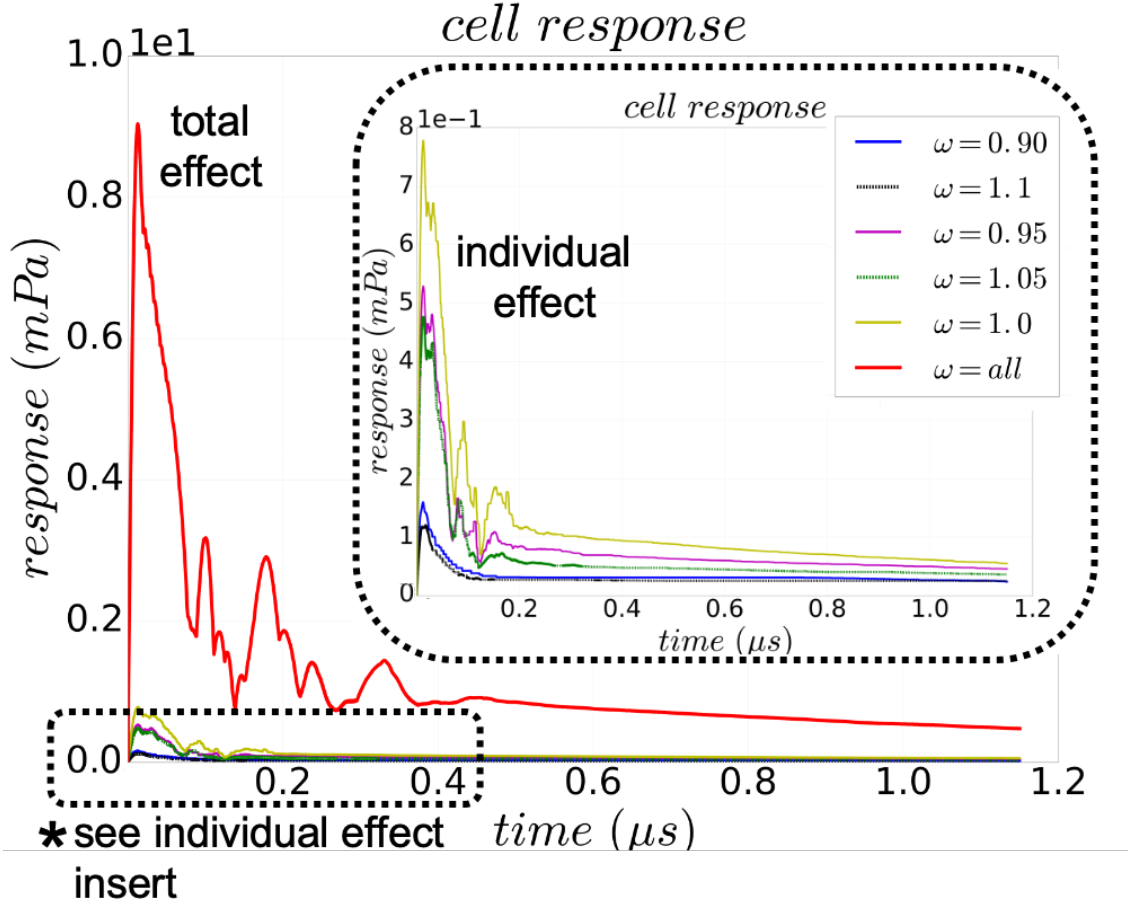
Temporal evolution of maximum elastic response magnitude for each frequency-amplitude coupling and from total effect. Peak cell frequency is in phase with source operating frequency with initial condition given by a phase of 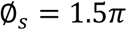.

**Figure 5.**
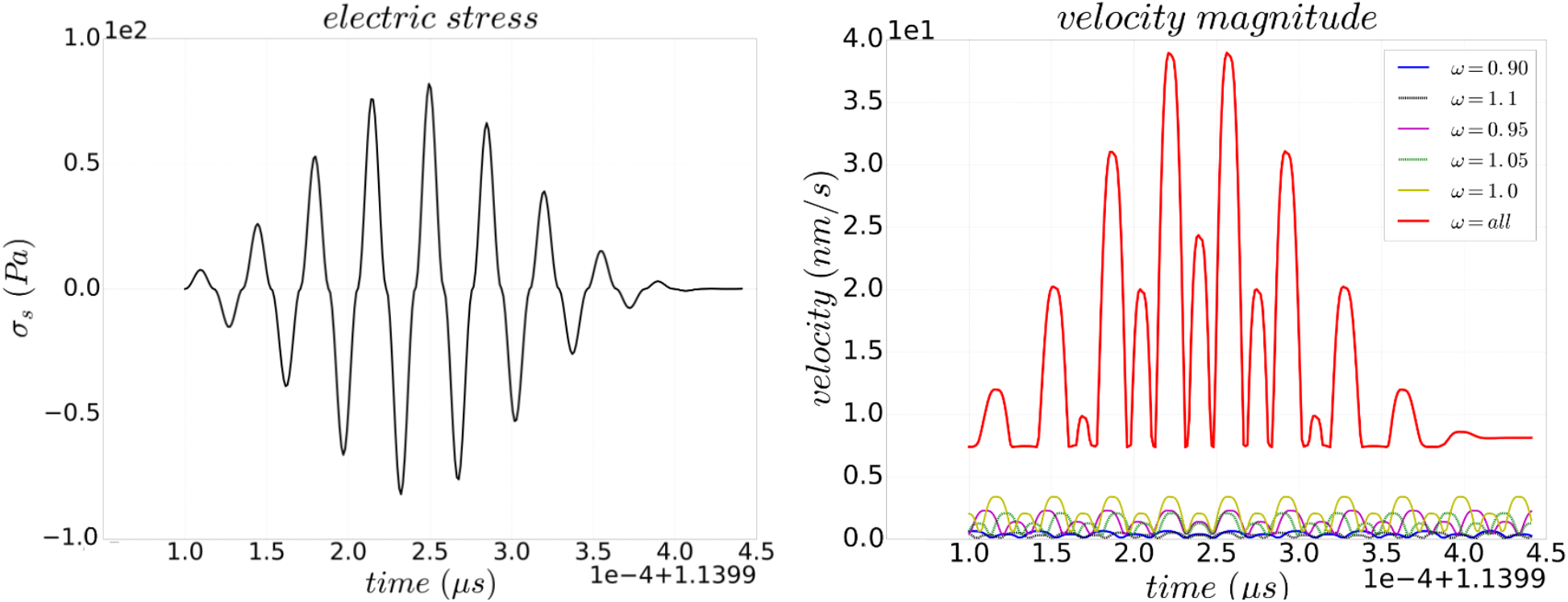
A complete stress cycle (left) and temporal evolution of maximum steady velocity (right) for each frequency-amplitude coupling and from total effect for each cycle. Peak cell frequency is in phase with source operating frequency with initial condition given by a phase of 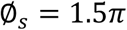.

The intensity and damping coefficient of oscillation can be obtained from response and cell velocity via equation (2.12). Figure 6 shows that maximum oscillation intensity occurs at resonance. For each trial, maximum damping occurs as the cell velocity approaches its minimum; so it follows that maximum intensity occurs as the cell velocity approaches its maximum with low damping. The effective damping coefficient, which is a contribution from the entire spectrum is lower than the individual contributions. Similarly, the total intensity is much higher than the individual contributions. These results highlight the importance of determining which source frequencies affect cell deformation since the total effect is not simply a sum of the individual contributions.

**Figure 6.**
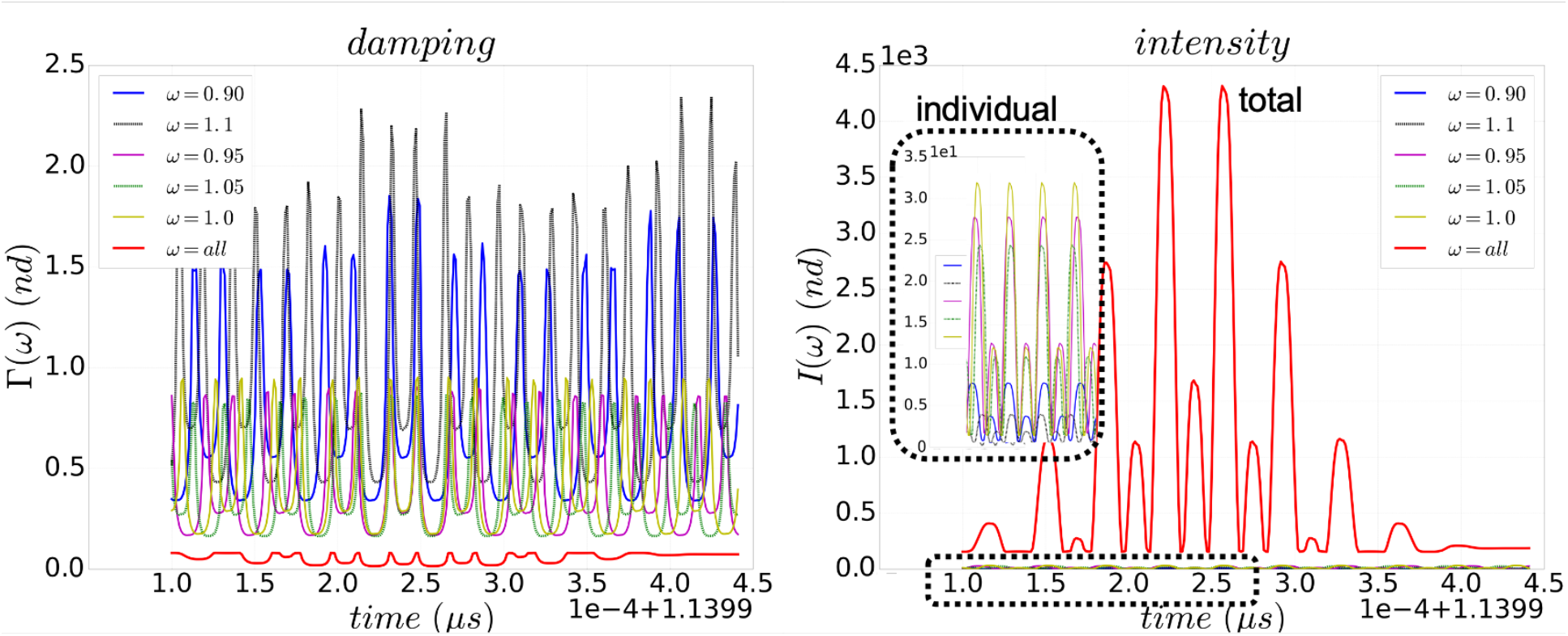
Damping and intensity of oscillation induced on the cell by individual frequency-amplitude couplings and by total effect. Peak cell frequency is in phase with source operating frequency with initial condition given by a phase of 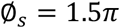. Results correspond with the electric stress shown in Figure 5.

Figure 7 shows the change in internal energy of the system due to the absorption of energy from the EM source minus the depletion via cell response. This figure shows that for each cycle at steady state, the maximum increase of internal energy is about 80 mJ/kg, with the accumulation of energy per cycle reaching almost 0.6 J/kg. The amount of energy expended through viscoelastic response is higher in the case of the total effect, and so is the energy delivered to the cell for the entire spectrum when compared to individual frequencies. To further understand the energy absorption by a typical cell, consider that the mass of typical micron-sized eukaryotic cells is in the nanogram range.^48^ Using this estimate, the maximum amount of energy that would be absorbed in one stress cycle would be in the nano-Joule range (0.6 J/kg x 1e-9 kg = 0.6 nJ).

**Figure 7.**
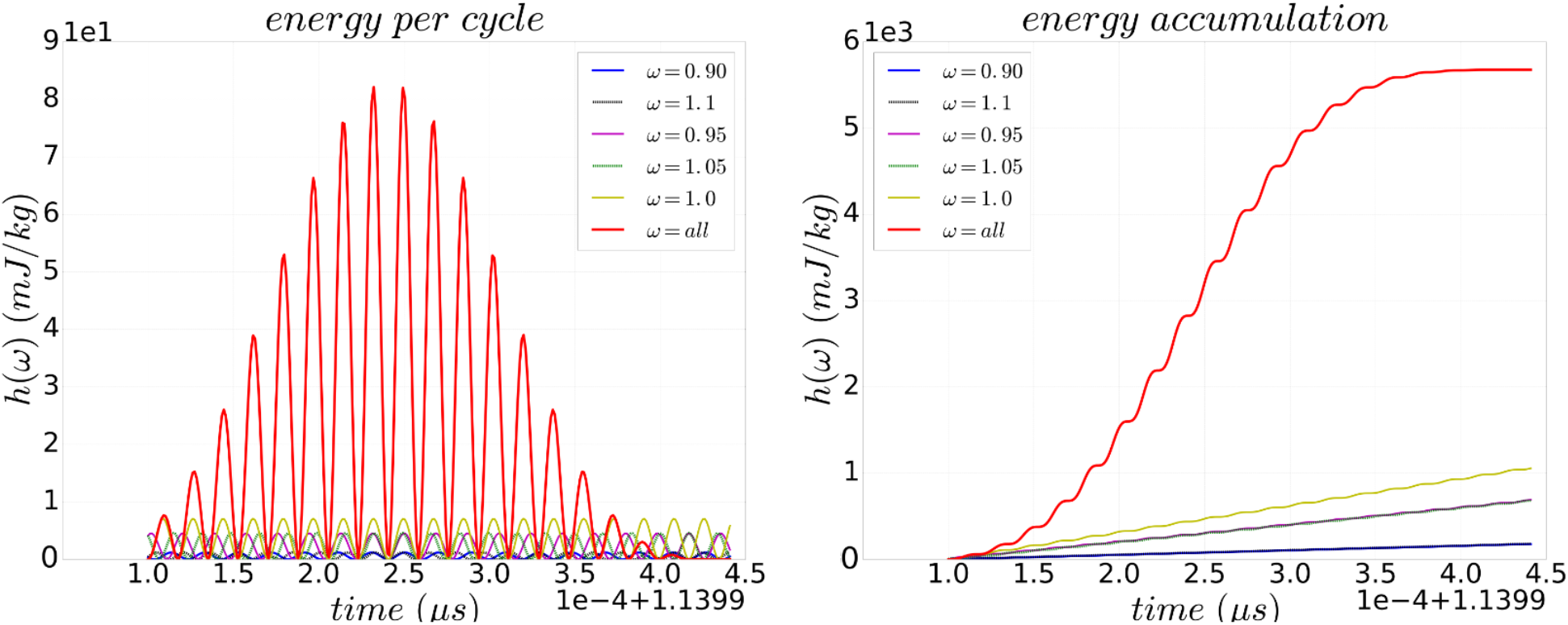
(left) Enthalpy change in the cell per cycle as a result of absorption from the EMF source and depletion from cell response. (right) enthalpy accumulation in cell from EMF source. Peak cell frequency is in phase with source operating frequency with initial condition given by a phase of 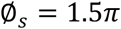.

Another calculation can be made to provide insight into the increase in temperature of the cell per cycle. Ahuja et al. estimated that at 25 degrees Celsius, the thermal conductivity of neuroblastoma tumor cells was 89 percent to that of water.^49^ Similarly, the thermal diffusivity of these cancer cells was 93 percent to that of water. Using these values, and assuming that the density of these cells is similar to the density of water, the specific heat of neuroblastoma cells can be estimated to be about 4 kJ/kg-K. Then if neuroblastoma cells accumulated 0.6 J/kg in one cycle, the increase in temperature would be small on the order of 15 micro-Kelvin per cycle. This result does not include energy expenditure from cell metabolic activity, which may occur in longer time scales so may not contribute in this calculation. The dissipation of heat by convection or diffusion may play important roles at these time scales, however. Further calculations must be done to understand how much energy absorbed per cycle is accumulated over longer time scales.

Spatial resolution of the cell as it is driven by the EM source provides additional insight into the effect of deformation. Figure 8 shows a snapshot of the deformed cell with an oscillating electric field in the x-direction as shown in Figure 1. The cell experiences the highest elastic expansion and compression in the direction perpendicular to the propagation of the EM field. This is consistent with the velocity field inside the cell, where divergent velocity vectors induce a restoring compression force, and convergent velocity vectors induce a corresponding expansion force. Video 1 provided in the supplementary materials shows an animation of the simulated velocity evolution during the steady state period with a delivered stress as shown in Figure 5. Figure 9 shows the dynamic pressure exerted by the fluid as a result of motion, as well as the maximum instantaneous energy accumulated in one stress cycle.

**Figure 8.**
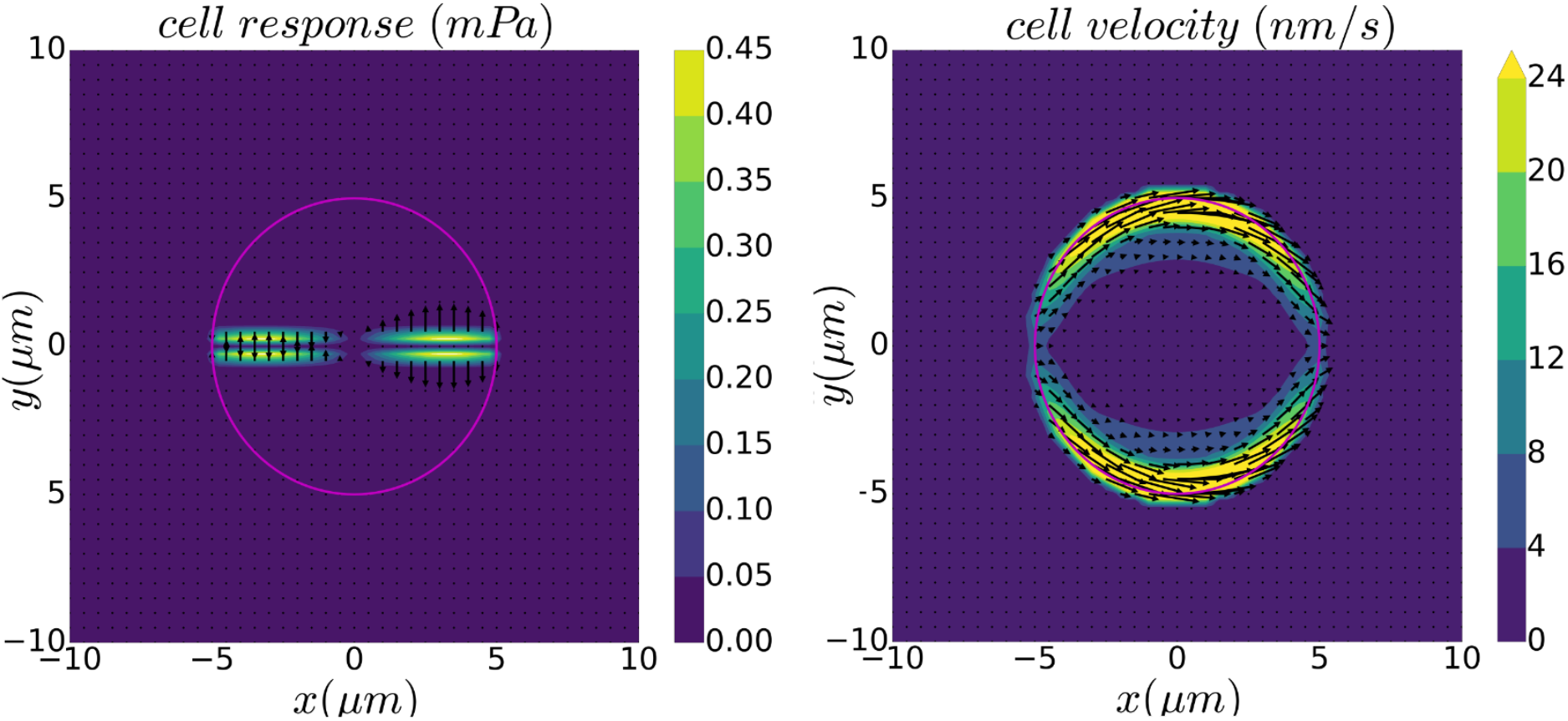
Spatial resolution of cell response (left) and velocity (right) from total effect. The plot corresponds to highest velocity induced by the electric stress cycle as shown in Figure 5. Maximum velocity is about 40 nm/sec. Peak cell frequency is in phase with source operating frequency with initial condition given by a phase of 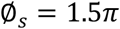. Circular outline represents the cell membrane.

**Figure 9.**
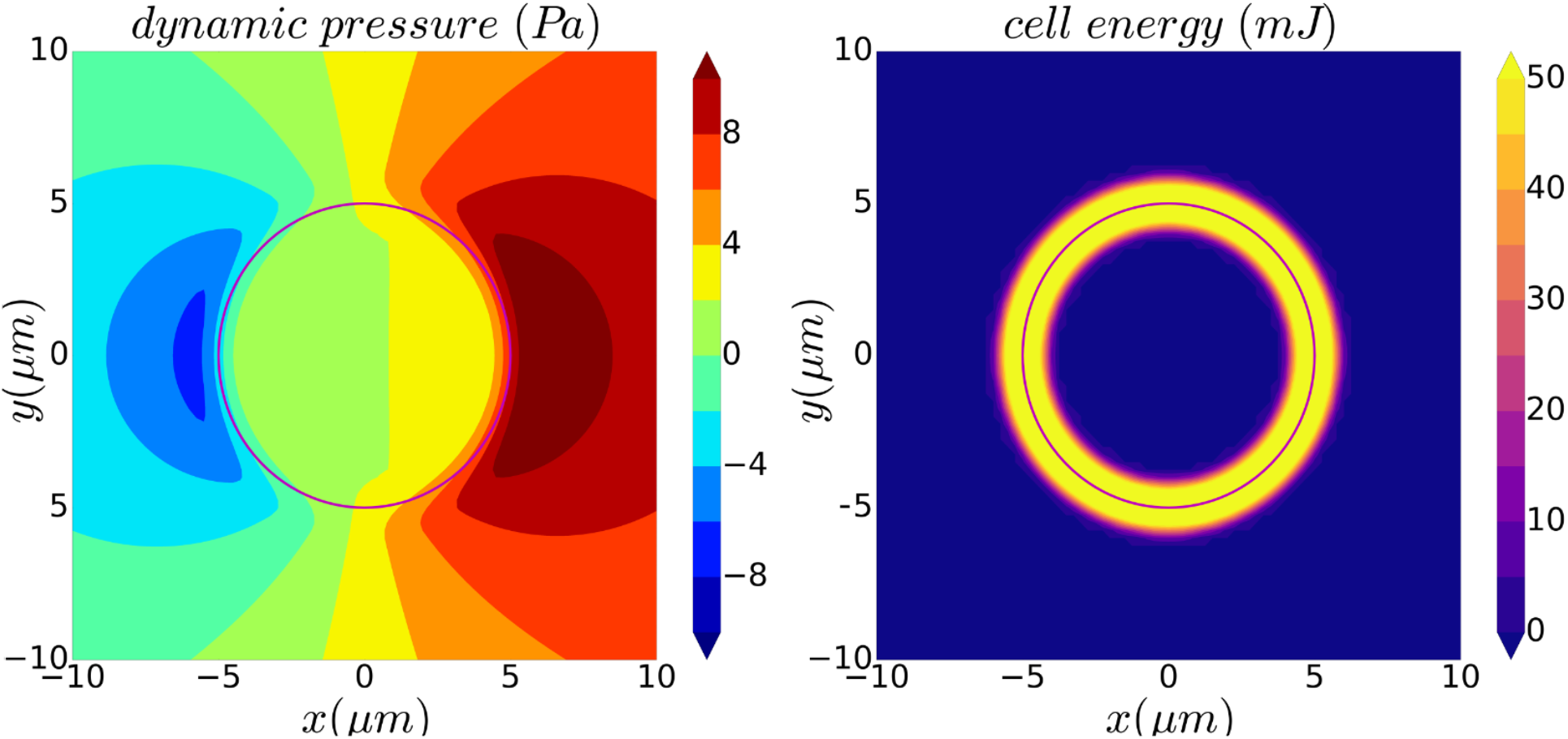
Spatial resolution of dynamic pressure (left) and energy (right) from total effect. The plot corresponds to highest pressure and accumulated energy caused by the electric stress cycle as shown in Figure 5. Peak cell frequency is in phase with source operating frequency with initial condition given by a phase of 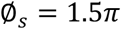. Circular outline represents the cell membrane.

### 3.2 Sensitivity of the model to source phase, elasticity, and source amplitudes

This section investigates the impact that several input parameters have on the transient and steady state solution to the cell deformation problem. First, the phase at which the source EM field begins to affect the cell defines the initial condition for the electric stress. Figure 10 shows two solutions to cell response and velocity magnitude for sources with the same energy density distribution but different incoming phases of 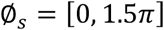. This result shows that the incoming phase of the EM source signal affects both the transient and steady state values of cell response and velocity. Figure 10 suggests that the source phase produces multiple solutions. However, the results are comparable in that the steady state values are of the same order of magnitude, and this means that the threshold necessary to induce some cell metabolic activity as a result of stress would be achieved regardless of the source phase. Ultimately, the amplitude of the source energy density will dictate the order of magnitude of cell response.

**Figure 10.**
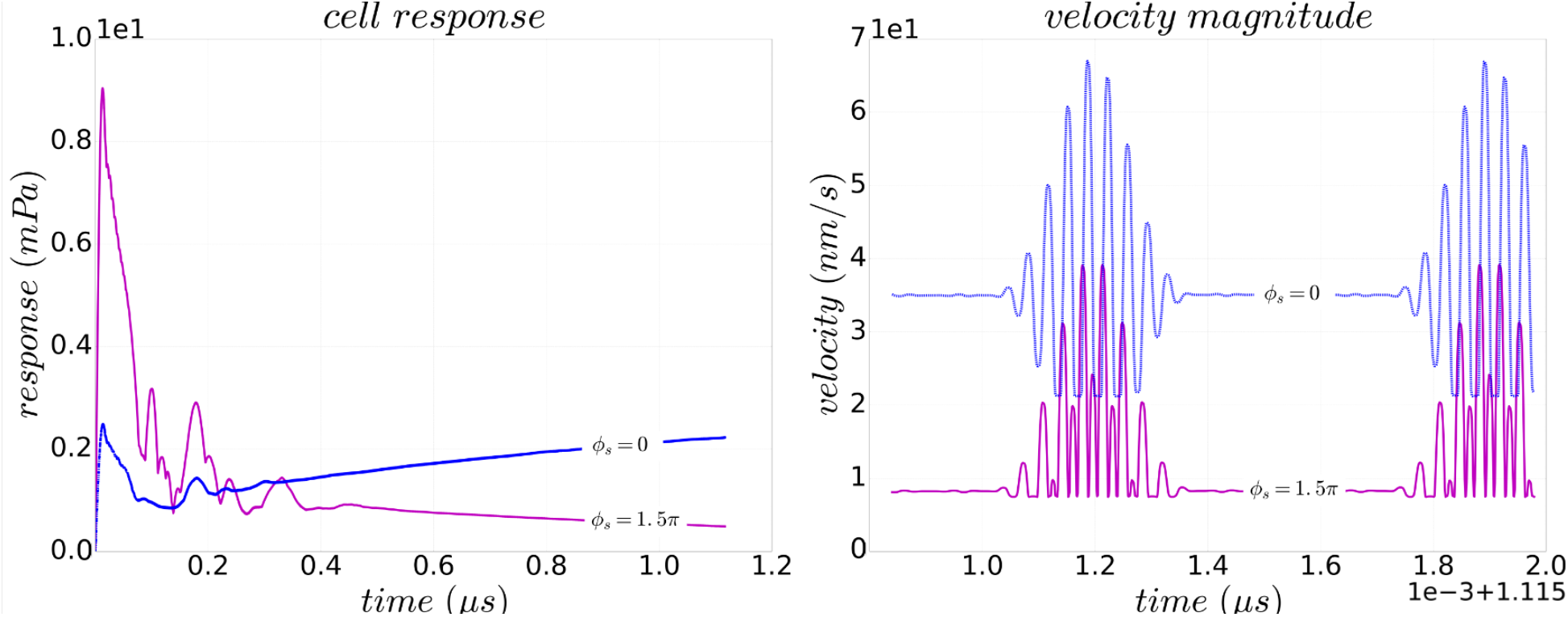
Multiple solutions to cell elastic response and velocity based on incoming source phase. Both trials have an EM source with all five frequency-amplitude couplings shown in Table 1.

For an EM source with a higher energy density, it is possible to induce a higher elastic response and even plastic deformation and rupture. For these limiting cases, absorption of energy would have to be bounded by the limits of the capacity of the cell to accumulate energy besides the electromechanical effect in deformation. This result is included in Appendix C of supplementary materials. As discussed in Section 1, a range of elastic moduli have been reported for both health and cancerous biological cells of different types, as well as for elastic properties of different components of cells. For the purpose of this study, a parametric test is included to demonstrate the effect of varying the effective elastic modulus of the cell structure (while leaving the Reynolds number constant). Figure 11 shows maximum cell response for three separate trials in which elastic modulus was varied. These results confirm that a higher elastic modulus scales the cell response force accordingly. The maximum cell oscillating velocity at steady state is affected by the elastic modulus of the cell; a higher elasticity yields a slightly lower velocity during a stress cycle. Damping coefficients differentiate the cases by showing that for elastic moduli lower than 10^7^ Pa, the system has minimal damping, with intensity of oscillation being different for each case.

**Figure 11.**
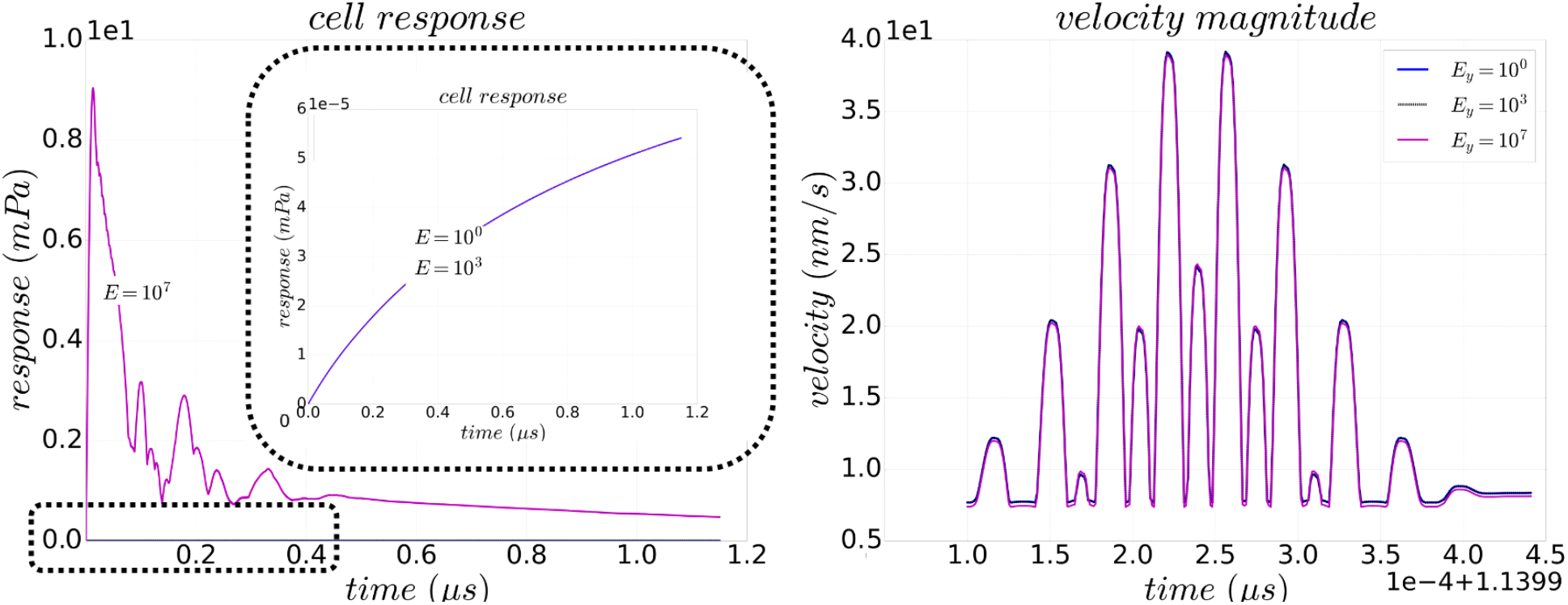
Comparison of cell response and oscillating velocity for different elastic moduli from total source effect. Peak cell frequency is in phase with source operating frequency with initial condition given by a phase of 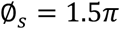. Results for velocity correspond with the electric stress shown in Figure 5.

**Figure 12.**
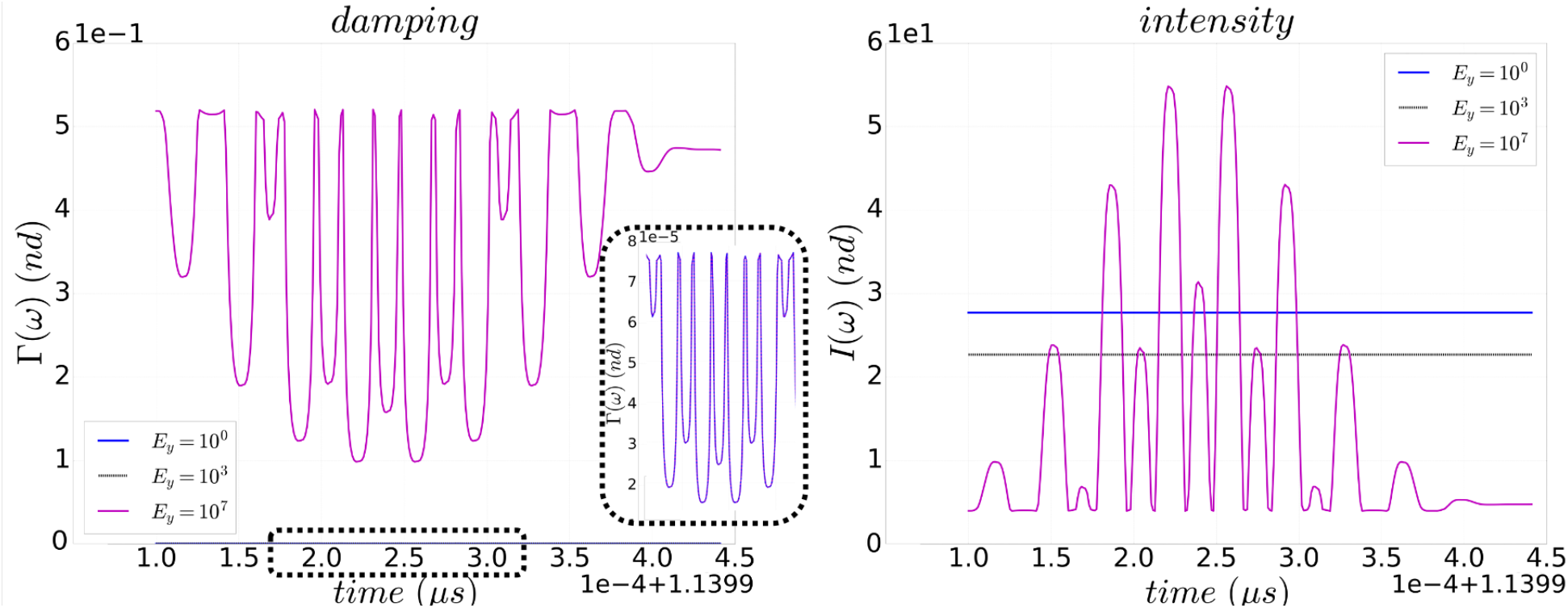
Damping and intensity of oscillation induced on the cell by total effect for varying elastic moduli. Peak cell frequency is in phase with source operating frequency with initial condition given by a phase of 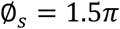. Results correspond with the electric stress shown in Figure 5.

## 4 Concluding Remarks

The model presented in this paper defines a biological cell as a structure susceptible to deformation with elastic response induced from external stresses. This approximation is consistent with existing experimental reports in which cells are deformed by mechanic stresses (e.g., parallel plate method) as well as electric stresses (e.g., optical tweezers), and for which elastic response and deformation were reported. In our work, the external stress is provided from an atmospheric plasma jet which delivers a spectrum of frequency-energy couplings within a narrow band centered around the frequency corresponding to the peak plasma density. We assume that the frequency at which the peak plasma density (and energy) occurs coincides with the eigenfrequency of the cell. While it may be practically difficult to create this scenario experimentally, it is certainly not impossible given that jet parameters which affect density can be manipulated. These include discharge voltage, gas flow rate, and humidity of background air. More importantly, in order to induce cell response, the jet does not need to deliver a resonant frequency-energy coupling, only a combination of near-resonant couplings defined by the spectrum of the source energy distribution function. In other words, the source frequency spectrum needs only to overlap the cell eigenfrequency to induce a response.

The model is limited by assuming that the cell’s elastic modulus is isotropic, and that the cell behaves as a highly viscous gel-like solid. The trends presented in this work are limited to a specific set of characteristic values. The choice of these values is informed by a literature review which suggests, in particular, a wide range of cell elastic moduli for biological cells. The characteristic cell density and viscosity were assumed to be those of water since the cells of interest mostly consist of water. The characteristic velocity of the cell is based on a formulation similar to that of sound speed in a medium, where it can be approximated as the square root of the ratio of elastic modulus to density. The nondimensionalization of the momentum equation yields an important number, here named frequency number (Nf), which appears in the source term and couples the driving frequency with the characteristic frequency of the cell. The frequency number is useful in comparing the time of each driving cycle relative to the behavior of the cell during this cycle. Physically, the implication of Nf = 1 is that the response of the entire cell is instantaneous to the applied stress, which would not be a common occurrence when the stress is applied locally on the membrane. A more probably range of Nf is greater than one, and it implies that the time to complete one characteristic driving stress cycle is smaller than the time it would take for the cell to fully respond to that cycle.

Results show that, when considering individual frequency-energy couplings, the maximum deformation effect occurs at the resonant frequency. Similarly, by approximating the source spectrum as five couplings acting simultaneously, the total deformation effect is greater than the sum of the individual contributions, which makes the solution nonlinear. Additionally, the phase of the source spectrum signal relative to the initial contact with the cell affects the transient and steady state behavior of deformation. This points to a scenario in which there are multiple solutions within the same order of magnitude for elastic response and velocity for the same experimental setup. Results show that higher cell response may be biased towards one half of the spectrum even if it is defined by a normal distribution. This dynamic should be resolved on a case-by-case basis as it depends on other variables, including the distribution shape of the energy source, the ratio of the driving to response frequencies, and the mode of oscillation of the cell.

At the time of this study, experimentally validating the predicted cell oscillatory behavior is difficult given the high driving frequency and very small predicted cell velocity. However, researchers may inform the design and calibration of measurement tools by an order of magnitude estimate. For example, given the conditions presented in this work, cell velocity is predicted to be in the nanometer per second range, with a response frequency in the mega Hertz range (note Nf = 452). Experimentalists may use this calculation to build an interferometer expected to measure a distance in the order of 10^-15^ meters, which is small yet remarkably within the resolution of existing interferometers. These estimations will vary from case to case, and the predicted velocity could be even higher given a more powerful source. To this end, a sensitivity analysis included in the supplementary materials shows that at larger stress levels than those considered in Section 3.1, the cell may reach and surpass its elastic limit to the point of rupture. These results are consistent with previous findings that biological cells can be sensitized or destroyed by EMF radiation.

A significant factor for further investigation is the amount of energy which the cell can accumulate over a long time scale without incurring thermal damage. Results show that the cell absorbs energy from the source, from which some is converted into kinetic energy as the source induces the cell to respond. In one stress cycle, it is estimated that the cell temperature may increase in the order of micro-Kelvin. If the energy absorbed per stress cycle is not mostly dissipated by diffusion, convection, or instantaneous metabolic activity, then the accumulation of energy may lead to the increase of energy to the degree that heat absorption would destroy the cell. In other words, addition of heat may lead to phase changes within the cytoskeleton which may be another source of cell rupture.

## 5 Author Contributions

LM and AD developed the numerical method and computational algorithm with supervision from MK and EB. LM ran the simulation and processed the output data. LL performed experimental measurements for emitted plasma jet power with supervision from MK. LM wrote the manuscript with contributions from AD, LL, EB, and MK.

## 6 Acknowledgments

This work was supported by the National Science Foundation, Grant number 1747760.

